# Engineering and optimization of the 2-phenylethylglucosinolate production in *Nicotiana benthamiana* by combining biosynthetic genes from *Barbarea vulgaris* and *Arabidopsis thaliana*

**DOI:** 10.1101/2020.05.12.090720

**Authors:** Cuiwei Wang, Christoph Crocoll, Niels Agerbirk, Barbara Ann Halkier

## Abstract

Among the glucosinolate (GLS) defense compounds characteristic of the Brassicales order, several have been shown to promote human health. This includes 2-phenylethylglucosinolate (2PE) derived from homophenylalanine (HPhe). In this study, we used transient expression in *Nicotiana benthamiana* to validate and characterize previously predicted key genes in the 2PE biosynthetic pathway from *Barbarea vulgaris* and demonstrate the feasibility of engineering 2PE production. We used genes from *B. vulgaris* and *Arabidopsis thaliana*, in which the biosynthesis of GLSs is predominantly derived from HPhe and dihomomethionine, respectively. The resulting GLS profiles partially mirrored GLS profiles in the gene donor plant, but in both cases the profiles in *N. benthamiana* were wider than in the native plants. We found that *Bv*BCAT4 is a more efficient entry enzyme for biosynthesis of both HPhe and dihomomethionine and that MAM1 enzymes determine the chain-elongated profile. Co-expression of the chain elongation pathway and *CYP79F6* from *B. vulgaris* with the remaining aliphatic GLS core pathway genes from *A. thaliana*, demonstrated the feasibility of engineering production of 2PE in *N. benthamiana*. Noticeably, the HPhe-converting enzyme *Bv*CYP79F6 in the core GLS pathway belongs to the CYP79F subfamily, a family believed to have substrate specificity towards chain-elongated methionine derivatives. Replacing the *B. vulgaris* chain elongation pathway with a chimeric pathway consisting of *Bv*BCAT4, *Bv*MAM1, *At*IPMI and *At*IPMDH1 resulted in an additional 2-fold increase in 2PE production, demonstrating that chimeric pathway with genes from different species can increase flux and boost production in an engineered pathway. Our study provides a novel approach to produce the important HPhe and 2PE in a heterologous host. Chimeric engineering of a complex biosynthetic pathway enabled detailed understanding of catalytic properties of individual enzymes - a prerequisite for understanding biochemical evolution - and with biotechnological and plant breeding potentials of new-to-nature gene combinations.

## 1. Introduction

Glucosinolates (GLSs) are amino acid-derived defense compounds characteristic of the Brassicales order (Halkier and Gershenzon, 2006). The diversity of GLSs are determined by the variety of precursor amino acids, chain elongation of precursor amino acids, secondary modifications of amino acid-derived side chains and decoration on the glucose moiety (Blažević et al., 2020). 2-Phenylethylglucosinolate (2PE), with the trivial name gluconasturtiin, possesses multiple biological functions due to *in situ* formation of its myrosinase-mediated hydrolysis product, 2-phenylethyl isothiocyanate (PEITC). Release of PEITC from 2PE can be used as safe and environment-friendly pesticide with strong deterrent or toxic effects on pests, e.g. juveniles of the nematode *Globodera rostochiensis* (Serra et al., 2002). 2PE and PEITC are also intermediates in the biosynthesis of novel phytoalexins with promising bioactivities (Pedras and To, 2018). Furthermore, PEITC released from 2PE has application in biofumigation and food preservation due to its anti-microbial properties (Dufour et al., 2015; Gimsing and Kirkegaard, 2009; Smith and Kirkegaard, 2002). From a health perspective, a number of studies suggest that PEITC is an effective cancer chemo-preventive and chemo-therapeutic agent, as evidenced by *in vitro* and animal studies (Mitsiogianni et al., 2019). However, PEITC in itself is reactive and cytotoxic in larger quantities, which suggests that it should be handled and administered as the precursor 2PE. These properties have primed a desire to increase levels and purity of 2PE produced in plants or engineer 2PE production in microbes.

For agricultural and industrial purposes, modification of GLS composition has been investigated using various biotechnological approaches and in various organisms (reviewed in Petersen et al., 2018). The organisms used span from native GLS-producing plants, as whole plants, cell or hairy root cultures, to heterologous hosts, such as tobacco and the microbes *Escherichia coli* and *Saccharomyces cerevisiae*. A prerequisite for implementation of those approaches is the understanding of the GLS biosynthesis. Production of 2PE through heterologous expression of proposed genes in the pathway was the ultimate test of our understanding of the pathway, and enabled investigating the interplay between individual genes.

In a biosynthesis context, GLSs are classified according to their amino acid precursor and side chain skeleton (Blažević et al., 2020). Those with aliphatic side chains, derived from aliphatic amino acids, are collectively known as aliphatic GLSs (derived from methionine, alanine, valine, leucine, isoleucine, and glutamate). Those with arylalkyl side chains, derived from aromatic amino acids, are usually discussed separately as benzenic GLSs (derived from phenylalanine and tyrosine) and indole GLSs (derived from tryptophan), respectively. The biosynthetic pathways of GLSs generally comprise two or three phases: chain elongation of selected precursor amino acids (only for a subset of GLSs), formation of the GLS core structure, and secondary modifications of the amino acid-derived side chain (Sønderby et al., 2010). The chemical structure of 2PE contains a common GLS moiety connected with a phenylethyl group as the side chain (**Supplementary Fig. S1**). The 2PE structure suggests biosynthesis from phenylalanine via one-cycle chain elongation to generate homophenylalanine (HPhe) that subsequently is converted by the core structure pathway to 2PE (**Supplementary Fig. S1**). Indeed, classical research confirmed this hypothesis by tracer experiments and detection of key intermediates in 2PE containing plants (Underhill, 1968), but the responsible genes could not be discovered at that time.

The genes in the methionine chain elongation pathway were first identified and characterized in *Arabidopsis thaliana*. First, methionine is deaminated by an aminotransferase (BCAT4) in the cytosol and the corresponding α-keto acid is imported by bile acid transporter 5 (BAT5) into the chloroplast. Here, it enters a cycle of three successive transformations: condensation with acetyl-CoA by methylthioalkylmalate synthase (MAM), isomerization by isopropylmalate isomerase (IPMI), and oxidative decarboxylation by isopropylmalate dehydrogenase (IPMDH1). The product of these three reactions is an α-ketoacid elongated by a single methylene group (-CH2-). Subsequently, the molecule can either be exported to the cytosol and transaminated, possibly by BCAT4 or other BCATs, or proceed through another round of chain elongation to synthesize higher homologs of methionine, such as dihomomethionine (DHM). MAM enzymes are critical in this respect as they determine the number of chain elongation cycles (Kroymann et al., 2003). In *A. thaliana*, MAM1 and MAM2 are involved in the biosynthesis of short-chain (C3-C5) GLSs and MAM3 in the biosynthesis of long-chain (C6-C8) GLSs (Benderoth et al., 2006; Textor et al., 2007, 2004; Field et al., 2004; Halkier and Gershenzon, 2006).

The core structure pathway is common for all GLS biosynthesis and contains seven enzymatic steps with the substrate-specific cytochrome P450s of the CYP79 family as the entry point (Sønderby et al., 2010). The CYP79 enzymes catalyze the conversion of precursor amino acids into oximes, and their substrate specificities are generally critical for the GLS products being generated. For example, *A. thaliana* CYP79F1 converts chain-elongated methionine derivatives (Chen et al., 2003) and *A. thaliana* CYP79A2 converts phenylalanine into the corresponding oxime (Wittstock and Halkier, 2000), although recent results suggest that CYP79s have a broader substrate specificity than realized *in planta* (Wang et al., 2020). The oximes are converted to the corresponding GLSs by downstream enzymes in the core structure pathway. Briefly, the oxime is oxidized by CYP83s to reactive nitrile oxides that are conjugated with glutathione by glutathione-S-transferases (GSTs). Cleavage by γ-glutamate peptidases (GGPs) forms *S*-alkylthiohydroximates, which are subsequently cleaved by C-S lyases to produce thiohydroximates that are *S*-glucosylated by glucosyltransferases, UGTs of the 74 family, to form desulfo-GLSs. Finally, desulfo-GLSs are sulfated by sulfotransferases (SOTs) to produce GLSs. Hitherto, in *A. thaliana*, the post-oxime enzymes in the core structure pathway have been divided into the so-called “aromatic” pathway consisting of CYP83B1, GSTF9, GGP1, SUR1, UGT74B1 and SOT16, and “aliphatic” pathway consisting of CYP83A1, GSTF11, GGP1, SUR1, UGT74C1 and SOT18 (Sønderby et al., 2010). Recently, we have shown that the post-oxime enzymes in both core structure pathways are capable of efficient biosynthesis of both aromatic and aliphatic GLSs (Wang et al., 2020). In the rest of this paper, the classical designations “aromatic” and “aliphatic” are still used for simplicity, although the current paper confirms that the designations are not adequate for describing the specificity of each group of core structure pathway enzymes.

2PE is one of the most common GLSs in plants of the Brassicaceae family, including roots of all the common *Brassica* vegetables (cabbages) (Van Dam et al., 2009), and the model plant *A. thaliana* (Wittstock and Halkier, 2000). It is particularly prominent in just a few species, including the aquatic herb watercress *(Nasturtium officinale)* (Gadamer, 1899; Agerbirk et al., 2014), and species in the *Barbarea* genus (Agerbirk and Olsen, 2011). A high-quality transcriptomics dataset has been obtained from G-type (pest-resistant) and P-type (pest-susceptible) *Barbarea vulgaris* R. Br., a wild cruciferous plant (Wei et al., 2013, Zhang et al., 2015). Recently, the dataset was mined specifically for predicted GLS biosynthetic genes based on homology searching with the genes known from *A. thaliana* (Liu et al., 2016). Roots of *B. vulgaris* are enriched in 2PE, but the transcriptome data were from leaves, where a hydroxyl derivative of 2PE rather than 2PE dominated. However, since 2PE is precursor for the hydroxylated 2PE derivative (van Leur et al., 2006; Liu et al., 2019), biosynthetic genes in 2PE biosynthesis was deduced to exist in *B. vulgaris* leaves, suggesting that the transcriptome data from leaves would be useful for gene discovery in an engineering context.

The lack of methionine-derived GLSs in *B. vulgaris* is an evolutionary mystery, since it is nested in a family and tribe characterized by this group of GLSs, such as dominance of methionine-derived GLSs in *A. thaliana* (Petersen et al., 2002). The evolutionary transition in a common ancestor of the genus *Barbarea* was enigmatic but appeared to be of a dual nature, both blocking biosynthesis of methionine-derived GLSs and promoting GLSs derived from phenylalanine (Liu et al., 2016; Byrne et al., 2017). Hence, experimental recombination of genes from the two biosynthetic processes of GLS could also shed light on this evolutionary question.

In this study, we demonstrate engineering of the production of HPhe from chain elongation biosynthetic genes of *B. vulgaris* and 2PE from a combination of genes of *B. vulgaris* and *A. thaliana* using *Nicotiana benthamiana* as heterologous host. Investigations of different combinations of genes from *B. vulgaris* and *A. thaliana* showed distinct effects on the profile of chain-elongated products as well as the flux through the 2PE pathway. The resulting GLS profiles were broader than in the source plants but showed clear effects of swapping critical genes among the biosyntheses.

## 2. Materials and methods

### 2.1. Candidate genes for the 2PE biosynthetic pathway from B. vulgaris

Putative coding sequences of genes of *B. vulgaris* in the phenylalanine chain elongation pathway and the first *CYP79F6* gene in the core structure pathway have been proposed from the *B. vulgaris* transcriptome data sets (supplementary material in Liu et al., 2016) based on homology searches with *A. thaliana* genes (*AtBCAT4* (AT3G19710), *AtMAM1* (AT5G23010), *AtIPMI-LSU1* (AT4G13430), *AtIPMI-SSU3* (AT3G58990), *AtIPMDH1* (AT5G14200) and *AtCYP79F1* (AT1G16410)). The CYP79F enzyme in *B. vulgaris* was named CYP79F6 according to the P450 nomenclature (Byrne et al., 2017), so this name is used here rather than the preliminary name (CYP79F1) used in Liu et al. (2016).

### 2.2. Generation and transformation of constructs

The gene sequences of *Bv*BCAT4 and *Bv*MAM1 were ordered as gBlocks from IDT (Integrated DNA Technologies Inc, USA). The gene sequences of *Bv*IPMI-LSU1, *Bv*IPMI-SSU3, *Bv*IPMDH1, *Bv*CYP79F6 were ordered from Twist Bioscience (San Francisco, USA). The gene sequences of *At*CYP79F1 and *At*CYP83A1 (AT4G13770) were PCR amplified from in-house plasmids (Mikkelsen et al., 2010). All DNA fragments of the genes were amplified by PCR with Phusion^®^ High Fidelity Polymerase (New England Biolabs, USA) and inserted into the vector pCAMBIA330035Su by USER cloning (Nour-Eldin et al., 2006). The NEB^®^ DH10B strain (New England Biolabs, #C3019H) was used to assemble and amplify the constructs. All the constructs used in this study are listed in **Table 1**.

**Table 1.** The constructs used in this study.

| Construct name | Expressed gene | Reference |
| --- | --- | --- |
| Construct 1 | *BvBCAT4* | This study |
| Construct 2 | *BvMAM1* | This study |
| Construct 3 | *BvIPMI-LSU1* | This study |
| Construct 4 | *BvIPMI-SSU3* | This study |
| Construct 5 | *BvIPMDH1* | This study |
| Construct 6 | *BvCYP79F6* | This study |
| Construct 7 | *AtCYP79F1* (AT1G16410) | This study |
| Construct 8 | *AtCYP83A1* (AT4G13770) | This study |
| p19 | *p19* | Crocoll et al., 2016b |
| *AtBCAT4* | *AtBCAT4* (AT3G19710) | Crocoll et al., 2016b |
| *AtMAM1* | *AtMAM1* (AT5G23010) | Crocoll et al., 2016b |
| *AtIPMI-LSU1* | *AtIPMI-LSU1* (AT4G13430) | Crocoll et al., 2016b |
| *AtIPMI-SSU3* | *AtIPMI-SSU3* (AT3G58990) | Crocoll et al., 2016b |
| *AtIPMDH1* | *AtIPMDH1* (AT5G14200) | Crocoll et al., 2016b |
| *AtBAT5* | *AtBAT5* (AT4G12030) | Crocoll et al., 2016b |
| GRN10 | *AtSOT18* (AT1G74090), *AtUGT74C1* (AT2G31790) and *AtSUR1* (AT2G20610) | Mikkelsen et al., 2010 |
| GRN11 | *AtGGP1* (AT4G30530) and *AtGSTF11* (AT3G03190) | Mikkelsen et al., 2010 |
| GRN12 | *AtCYP83A1* (AT4G13770) and *AtCYP79F1* (AT1G16410) | Mikkelsen et al., 2010 |
| ORF1+GGP1 | *AtCYP83B1* (AT4G31500), *AtCYP79A2* (AT5G05260) and *AtGGP1* (AT4G30530) | Geu-Flores et al., 2009 |
| ORF2 | *AtSOT16* (AT1G74100), *AtUGT74B1* (AT1G24100) and *AtSUR1* (AT2G20610) | Geu-Flores et al., 2009 |
| *AtAPK2* | *AtAPK2* (AT4G39940) | Møldrup et al., 2011 |

The constructs were transformed into *Agrobacterium tumefaciens* strain pGV3850 by electroporation (2 mm cuvette, 2.5 kV, 400 Ω, and 25 μF) in a Bio-Rad GenePulser (Bio-Rad, Hercules, CA, USA). Cells were incubated at 28°C for 2 h after 200 μl LB media was added. The culture was plated on LB agar plates containing 30 μg/ml rifampicin and 50 μg/ml kanamycin and incubated at 28°C for 3 days. Colony PCR was used to confirm the constructs in the strain. The primers used for cloning and colony PCR in this study were listed in **Supplementary Table S1**.

### 2.3. Transient expression of candidate genes in N. benthamiana

Strains of *A. tumefaciens* harboring different constructs were grown in YEP media containing 30 μg/ml rifampicin and 50 μg/ml kanamycin at 28°C and 170 rpm overnight. The cells were harvested by centrifugation at 4000 × *g* for 10 min at room temperature. Subsequently, the pellets were resuspended in infiltration buffer (10 mM MES, 10 mM MgCl_2_, pH 5.6) with 100 μM acetosyringone (3,5-dimethoxy-4-hydroxyacetophenone, Sigma-Aldrich, Steinheim, Germany) and slightly shaken at room temperature for 1-3 hours. The OD_600_ for each culture was measured and infiltration buffer was added to adjust to an OD_600_ ≈ 0.2. Equal volumes of each infiltration buffer containing individual constructs were mixed according to the experimental design. Combinations with fewer than seven constructs were adjusted with infiltration buffer, resulting in OD_600_ ≈ 0.029 for each individual construct. In the combinations with entire GLS biosynthetic pathways, infiltration buffer was added to combinations with fewer than 12 constructs to adjust each individual construct to OD_600_ ≈ 0.017. The silencing suppressor p19 was added to decrease silencing effects in all experiments (Voinnet et al., 2003). Two or three leaves of each *N. benthamiana* plant (around 4-week-old with four to six leaves stage) were infiltrated with mixed cultures of the different combinations.

### 2.4. Harvest of plant material and metabolite extraction

5-days after infiltration, four leaf disks of 1 cm diameter were harvested from each infiltrated leaf and weighed. The total metabolites were extracted using 85% methanol and the extract was diluted 10 times (1 vol. extract:9 vol. stock solution) with a stock solution containing 10 μg/ml ^13^C-, ^15^N-labelled amino acids (Algal amino acids ^13^C, ^15^N, Isotec, Miamisburg, US) and 2 μM sinigrin (PhytoLab, Vestenbergsgreuth, Germany). The diluted samples were filtered (Durapore^®^ 0.22 μm PVDF filters, Merck Millipore, Tullagreen, Ireland) and used directly for LC-MS analysis.

### 2.5. Analysis by LC-MS

Amino acids, chain-elongated derivatives and GLSs in the diluted extracts were directly analyzed by LC-MS as previously described (Mirza et al., 2016) with changes as detailed below. Chromatography was performed using an Advance UHPLC system (Bruker, Bremen, Germany) and a Zorbax Eclipse XDB-C18 column (100 x 3.0 mm, 1.8 μm, Agilent Technologies, Germany). Mobile phase A was water supplied with formic acid (0.05% (v/v), mobile phase B was acetonitrile supplied with formic acid (0.05% (v/v). The elution profile was: 0-1.2 min, isocratic 3% B; 1.2-4.3 min, linear gradient 3-65% B; 4.3-4.4 min, linear gradient 65-100% B; 4.4-4.9 min, isocratic 100% B, 4.9-5.0 min, linear gradient 100-3% B; 5.0-6.0 min, isocratic 3% B. Mobile phase flow rate was 500 μL/min and column temperature was maintained at 40°C. An EVOQ Elite TripleQuadrupole mass spectrometer with an electrospray ionization source (ESI) (Bruker, Bremen, Germany) was coupled to the liquid chromatography. The ion spray voltage was maintained at 3000 V or −4000 V in positive or negative ionization mode, respectively. Cone temperature was set to 350°C, cone gas flow to 20 psi, heated probe temperature to 400°C, probe gas flow to 50 psi, nebulizing gas to 60 psi, and collision gas to 1.6 mTorr. Nitrogen was used as both cone gas and nebulizing gas and argon as collision gas. Multiple reaction monitoring (MRM) was used to monitor transitions of analyte parent ions to product ions: MRMs for amino acids were chosen as previously described (Jander et al., 2004); MRMs for chain-elongated amino acids were chosen as described in (Mirza et al., 2016; Petersen et al., 2019b); MRMs for 2PE, 4-(methylthio)butylglucosinolate (4MTB), 4-(methylsulfinyl)butylglucosinolate (4MSB), 3-(methylsulfinyl)propylglucosinolate (3MSP) and benzylglucosinolate (BGLS) were chosen as previously reported (Crocoll et al., 2016a). Both Q1 and Q3 quadrupoles were maintained at unit resolution. Bruker MS Workstation software (Version 8.2.1, Bruker, Bremen, Germany) was used for data acquisition and processing.

For conclusive identification of homologs of leucine (that would be isomeric with homologs of isoleucine and higher homologs of alanine), we used homoleucine (HL) as reference standard as described in Mirza et al. (2016). Leucine homologs with longer side chain were therefore assumed to also be derived from leucine rather than isoleucine. Leucine and isoleucine are baseline separated with the column used for liquid chromatography and therefore it was concluded that homologs would be separated.

A mixture of ^15^N-, ^13^C-labeled amino acids was used as internal standard for quantification of amino acids and chain-elongated amino acids. Sinigrin was used as the internal standard for quantification of 2PE, 4MTB, 4MSB, 3MSP and BGLS. Response factors were calculated by dilution series of the respective analytes, based on matching ionization mode of the target analytes, negative ionization mode for GLSs and positive ionization mode for amino acids and chain-elongated amino acids. Due to matrix effect from the extract on the quantification of the analytes, the correction factors were calculated into the response factors in **Supplementary Table S2**. For analytes multiple transitions were monitored, the transition used for quantification is marked as quantifier (Qt). Further information on transitions and collision energies can be found in **Supplementary Table S2**.

## 3. Results

### 3.1. *Amino acid product profiles upon expression of the B. vulgaris* and *A. thaliana chain elongation pathways in N. benthamiana*

By homology-based searches in the transcriptome dataset of *B. vulgaris* using *A. thaliana* orthologs as reference, five candidate genes - *BvBCAT4, BvMAM1, BvIPMI-LSU1, BvIPMI-SSU3* and *BvIPMDH1* - in the phenylalanine chain elongation pathway have been proposed (Liu et al., 2016). The similarity of amino acid sequence between orthologs of *BCAT4* and *MAM1* from *B. vulgaris* and *A. thaliana* are around 80%, while orthologs of IPMI-LSU1, IPMI-SSU3 and IPMDH1 share identity above 87% (**Table 2**).

**Table 2.** Predicted genes in the phenylalanine chain elongation pathway from *B. vulgaris*.

| Candidate genes | Length (bp) | Genotype | CDS similarity with *A. thaliana* orthologs (%) | AA similarity with *A. thaliana* orthologs (%) |
| --- | --- | --- | --- | --- |
| *BvBCAT4* | 1065 | G and P | 86.17 | 79.83 |
| *BvMAM1* | 1512 | P | 86.57 | 80.47 |
| *BvIPMI-LSU1* | 1539 | P | 92.46 | 96.67 |
| *BvIPMI-SSU3* | 762 | G | 87.55 | 87.40 |
| *BvIPMDH1* | 1218 | P | 91.28 | 92.93 |

CDS represents gene coding sequence and AA represents amino acid sequence.

To test if we could produce HPhe, we transiently expressed the five *B. vulgaris* candidate genes for phenylalanine chain elongation in tobacco and analyzed the resulting levels of chain elongated amino acids. The following discussion is based on mean levels unless otherwise specified. Bile acid transporter 5 (BAT5) from *A. thaliana* was included to facilitate transport of α-keto acids between the chloroplast and cytosol (Crocoll et al., 2016b). We named this combination of chain elongation pathway enzymes C1 (**Fig. 1A**). We found that expression of the *B. vulgaris* combination C1 resulted in HPhe levels at 15 nmol/g fw (**Fig. 1B** and **Supplementary Table S3**), which confirms that the genes are able to chain elongate phenylalanine. Unexpectedly, the chain-elongated methionine derivative DHM - the precursor amino acid for 4MSB - accumulated at higher levels, 24 nmol/g fw (**Fig. 1C** and **Supplementary Table S3**). Detailed product profile analysis revealed that the *B. vulgaris* enzymes produced additional homologs of methionine and also chain elongated leucine, resulting in accumulation of homomethionine (HM), trihomomethionine (THM), homoleucine (HL) and dihomoleucine (DHL) (**Supplementary Fig. S2**). These results show that the *B. vulgaris* enzymes are capable of chain elongating phenylalanine as well as methionine and leucine when transiently expressed in tobacco, although in the native plant, *B. vulgaris*, GLSs derived from aliphatic amino acids do not accumulate (Byrne et al. 2017).

**Fig. 1.**
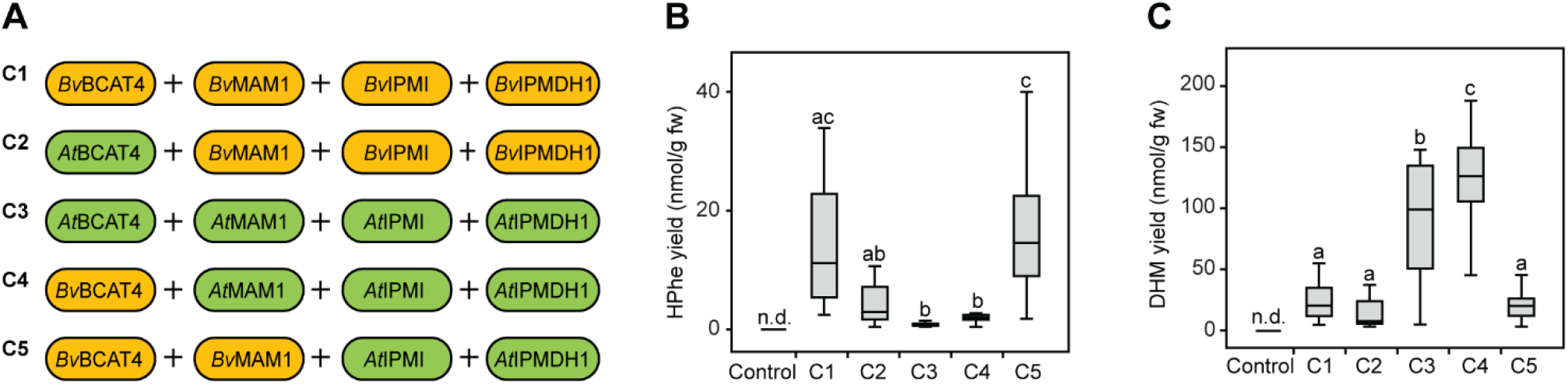
Production of HPhe and DHM in *N. benthamiana* upon transient expression of different chain elongation combinations. **A)** Scheme of chain elongation combinations containing different orthologs of BCAT4, MAM1, IPMI and IPMDH1. The *A. thaliana* transporter BAT5 is included in each combination. **B)** HPhe yields. **C)** DHM yields. Control plants were infiltrated with p19 and empty plasmid. Data of each box represent 16 biological replicates in nanomole per gram fresh weight. Data in **B** and **C** are box-and-whisker representations indicating the 10th (lower whisker), 25th (base of box), 75th (top of box) and 90th (top whisker) percentiles. The line within the box is the median and outliers are not shown. Exact values are listed in **Supplementary Table S3**. n.d. represents not detected. Letters represent significant differences by pairwise comparisons based on Tukey HSD test (*p* < 0.05) and *p*-values are listed in **Supplementary Table S4**. Data labelled with different letters are significantly different and the labels containing the same letter mean no significant difference.

For comparison, we transiently expressed in tobacco the *A. thaliana* chain elongation pathway including *At*BCAT4, *At*MAM1, *At*IPMI (consisting of *At*LSU1 and *At*SSU3) and *At*IPMDH1, named C3 (**Fig. 1A**). This resulted in the accumulation of DHM and other derivatives of methionine and leucine (HM, THM, HL and DHL) as well as HPhe (**Fig. 1B** and **C**, and **Supplementary Fig. S2**), which is consistent with previous results obtained in *E. coli* (Petersen et al., 2019b). The *B. vulgaris* combination, C1, produced 17-fold higher HPhe levels than the *A. thaliana* combination, C3, whereas the *A. thaliana* combination produced 3.9-fold higher DHM levels than *B. vulgaris* combination (**Fig. 1B** and **C**, and **Supplementary Table S3**). The DHM level in combination C3 was much higher (92 nmol/g fw) than the HPhe level in combination C1 (15 nmol/g fw). This difference suggests that the *A. thaliana* chain elongation machinery is overall more efficient than the *B. vulgaris* machinery in forming high levels of the relevant chain elongated products when expressed in tobacco cells. Moreover, these product profiles are consistent with *B. vulgaris* mainly producing phenylalanine-derived GLSs and *A. thaliana* predominantly producing methionine-derived GLSs, although the product profile from each sets of genes was much less specific than seen in the native plant. In particular, high levels of THM as well as HL and DHL observed with both sets of genes (**Supplementary Fig. S2**) were in stark contrast to GLS profiles in the native plants.

### 3.2. BCAT4 and MAM1 are critical for the profile of chain-elongated products

BCAT4 is the first enzyme in the chain elongation pathway. To investigate the effect of the substrate specificity of the two *BCAT4* orthologs on production of chain-elongated derivatives, we generated the chimeric combination C2 containing *At*BCAT4 and the remaining enzymes from *B. vulgaris* (**Fig. 1A**). Likewise, we constructed the combination C4 harboring *Bv*BCAT4 and the remaining enzymes from *A. thaliana* (**Fig. 1A**).

Replacing the first *B. vulgaris* enzyme (BCAT4) with the *A. thaliana* homolog (combination C2) resulted in a tendency for lower production of HPhe when compared to combination C1 (all *B. vulgaris* enzymes) (*p* = 0.246 shown in **Supplementary Table S4**), but very similar levels were observed for production of other chain-elongated derivatives (**Fig. 1B** and **C**, **Supplementary Fig. S2**). The result suggests that *Bv*BCAT4 has higher catalytic efficiency towards phenylalanine than *At*BCAT4, although the effect was not statistically significant. Moreover, expression of the reciprocal combination C4 (all *A. thaliana* enzymes except BCAT4 from *B. vulgaris)* resulted in a statistically significant increase for accumulation of DHM and leucine-derived products, but similarly low levels of HPhe as C3 (**Fig. 1B** and **C,** and **Supplementary Fig. S2**). The result suggests that *Bv*BCAT4 has higher catalytic efficiency towards methionine and leucine than *At*BCAT4. Overall, *Bv*BCAT4 seemed to be a more efficient entry enzyme than *At*BCAT4 in producing either HPhe or DHM depending on the remaining enzymes in chain elongation pathway. However, a role of *Bv*BCAT4 in determining the profile of the chain-elongated amino acids in *B. vulgaris* was not evident.

Although BCAT4 is the chain elongation entry point, we found by comparing combinations C1 and C4 that *Bv*BCAT4 does not determine the product profile of the full set of *B. vulgaris* enzymes (C1) (**Fig. 1B** and **C**). Hence, we substituted *At*MAM1 with *Bv*MAM1 in combination C4 to generate C5 (**Fig. 1A**) to investigate whether *Bv*MAM1 could restore HPhe production and the product profile of C1. Indeed, co-expression of *Bv*MAM1 together with *Bv*BCAT4 restored HPhe production with a level of 20 nmol/g fw, equally high to the production level from *B. vulgaris* chain elongation pathway C1 (**Fig. 1B** and **Supplementary Table S3**). This result shows that *Bv*MAM1 is critical for high HPhe production. The chain-elongated derivatives from methionine and leucine also accumulated to similar levels as for C1 (**Fig. 1C** and **Supplementary Fig. S2**). The results demonstrate that the MAM1 enzymes determine the product profile. Furthermore, substitution of *At*MAM1 by *Bv*MAM1 decreased the accumulation of DHM, HM, HL and DHL compared to C4 (**Fig. 1C** and **Supplementary Fig. S2**). This decrease shows that *Bv*MAM1 displays a lower catalytic rate with the aliphatic amino acids compared to *At*MAM1. The combinations containing *At*MAM1 (C3 and C4) produced tiny amounts of HPhe but significantly higher levels of methionine- and leucine-derived chain-elongated products except THM (**Fig. 1B** and **C** and **Supplementary Fig. S2**). Overall, the results show that the catalytic properties of *Bv*MAM1 and *At*MAM1 are major determinants of the product profiles of the two chain elongation pathways when expressed in tobacco.

### 3.3. The aliphatic core structure pathway converts HPhe to 2PE

In the core structure pathway - the second stage of 2PE biosynthesis - HPhe is converted to 2PE. To test which core structure pathway is suitable for this process, we first co-expressed in tobacco the chain elongation combination C1 with, respectively, the aromatic (Geu-Flores et al., 2009) or aliphatic core structure pathways (Mikkelsen et al., 2010) from *A. thaliana* (**Fig. 3A**). The GSTF9 from the aromatic core structure pathway (and corresponding to enzyme GSTF11 in the aliphatic core structure pathway) was not included, but this deficiency has been shown not to affect the functionality of the pathway in tobacco (Geu-Flores et al., 2009). In addition, we co-expressed the APS kinase APK2 from *A. thaliana* (**Fig. 3A**) since it has been reported to be critical for efficient regeneration of the co-factor PAPS in the final sulfotransferase step that converts desulfo-GLSs to intact GLSs (Møldrup et al., 2011). The chemical structures of the GLSs detected in this study and their amino acid precursors and chain-elongated intermediates are shown in **Fig. 2**. We found that co-expression of the aliphatic and not the aromatic core structure pathway produced 2PE (**Fig. 3B**). This result shows that *At*CYP79F1 has substrate specificity towards HPhe (in addition to homologs of methionine), which has not been reported previously. Additionally, we also detected HM-derived 3MSP, DHM-derived 4MTB and 4MSB when co-expressing the aliphatic core structure pathway and chain elongation combination C1 (**Supplementary Fig. S3A**). The accumulation of 3MSP and 4MSB shows that even in the absence of the *S*-oxygenating FMO_GS-OX1_ enzyme methylthio groups are partially oxidized to methylsulfinyl group by endogenous enzymes as previously observed (Mikkelsen et al., 2010). This product profile shows that *At*CYP79F1 takes HM and DHM as substrate, which is consistent with our previous study (Wang et al., 2020). However, accumulation of HL and DHL-derived GLSs is not observed in the current study, which is in contrast to our previous result (Wang et al., 2020). A possible explanation could be that the *A. thaliana* combination C3 producing higher level of HL and DHL was used in the previous study and the *B. vulgaris* set C1 producing small amounts of HL and DHL was used in the current study (**Supplementary Fig. S2C** and **D**). Co-expression of the aromatic core structure pathway resulted in a high level of BGLS due to the presence of the phenylalanine-specific CYP79A2 in the construct (**Supplementary Fig. S3B**), showing that the aromatic core structure pathway is functionally expressed. Lack of 2PE production with this construct is in agreement with the known specificity of *At*CYP79A2 for phenylalanine and not HPhe when expressed in *E. coli* (Wittstock and Halkier, 2000). However, lack of 2PE production with the aromatic pathway does not exclude that the post-oxime enzymes in the aromatic pathway may be suitable for conversion of HPhe to 2PE. In summary, we have engineered the production of 2PE by combining the five-gene chain elongation pathway from *B. vulgaris* and the aliphatic core structure pathway from *A. thaliana*. We have furthermore shown that both sets of expressed biosynthetic genes are much less specific than in the source plants.

**Fig. 2.**
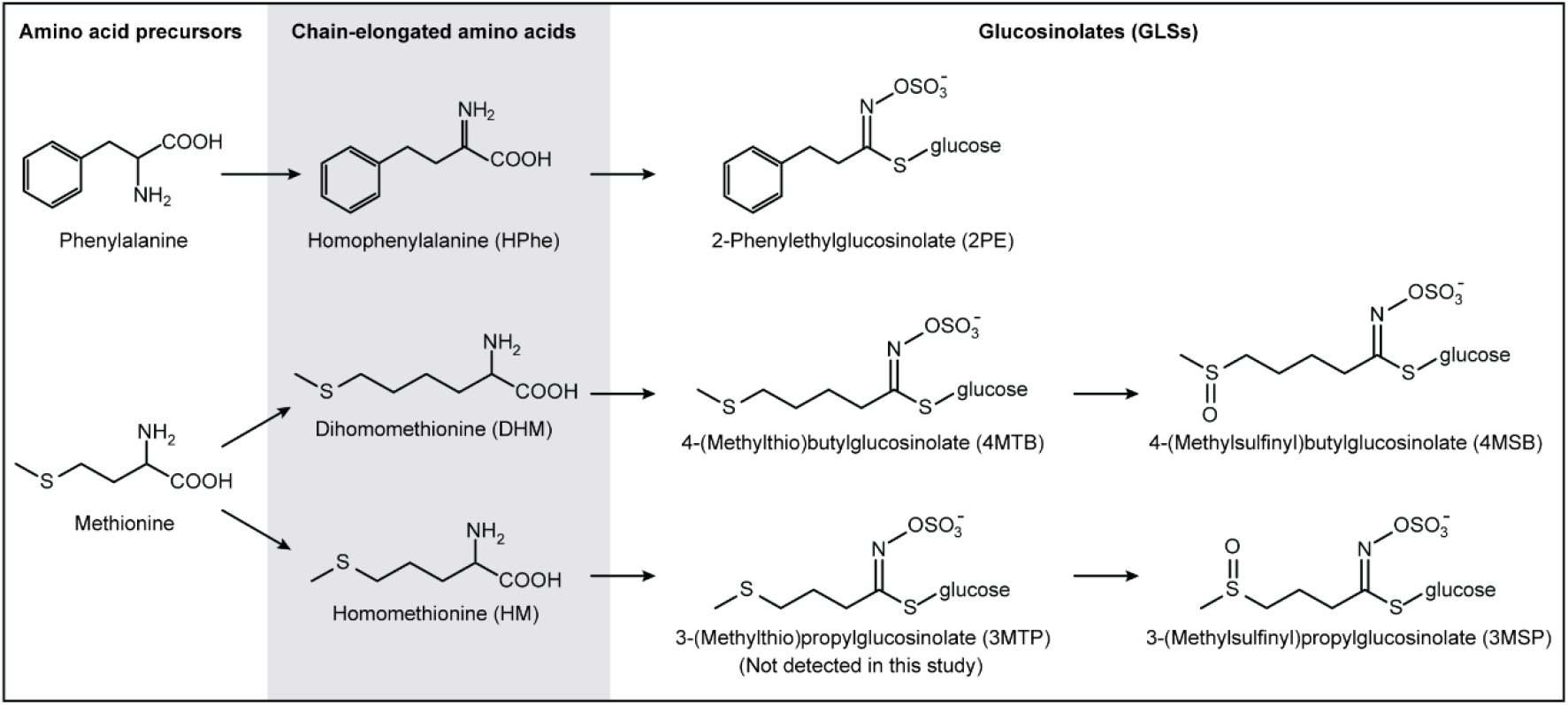
The chemical structures and biosynthetic relationships of amino acids and GLSs key to this study. 3-(Methylthio)propylglucosinolate (3MTP), the precursor of 3-(methylsulfinyl)propylglucosinolate (3MSP) was not detected in this study.

### 3.4. A chimeric chain elongation pathway enhanced 2PE production

Towards improving 2PE production in tobacco, we first combined the highest HPhe-producing chain elongation pathways, i.e. the *B. vulgaris* combination C1 and the chimeric combination C5, with the *A. thaliana* “aliphatic” core structure pathway (**Fig. 3C**), from now on called the core structure pathway. Although C1 and C5 produced similar high levels of HPhe (**Fig. 1B**), we found that co-expression with C5 resulted in an approximately 2-fold increase of 2PE production compared to co-expression of C1 (**Fig. 3D** and **Supplementary Table S3**). Similarly, the total yield of DHM-derived 4MTB and 4MSB was boosted by 2-fold when using the chimeric combination C5 (**Fig. 3E** and **Supplementary Table S3**). This suggests that the chimeric C5 pathway exhibits a higher flux of all intermediates, including HPhe and DHM than the *B. vulgaris* combination C1, when co-expressed with the core structure pathway that converts the chain-elongated amino acids to the corresponding GLSs. Despite the similar levels of HPhe and DHM observed when C1 and C5 were expressed alone, the 2PE yield from the complete GLS pathway (21 and 40 nmol/g fw, respectively) was significantly higher than the total yield of 4MTB and 4MSB (11 and 22 nmol/g fw, respectively) (**Supplementary Table S3**). This difference shows that co-expression of the core structure pathway has impact on the flux through the chain elongation pathway and suggests that the core structure pathway has a pull effect on 2PE accumulation, and to a relatively lower degree on the formation of DHM-derived 4MTB and 4MSB.

**Fig. 3.**
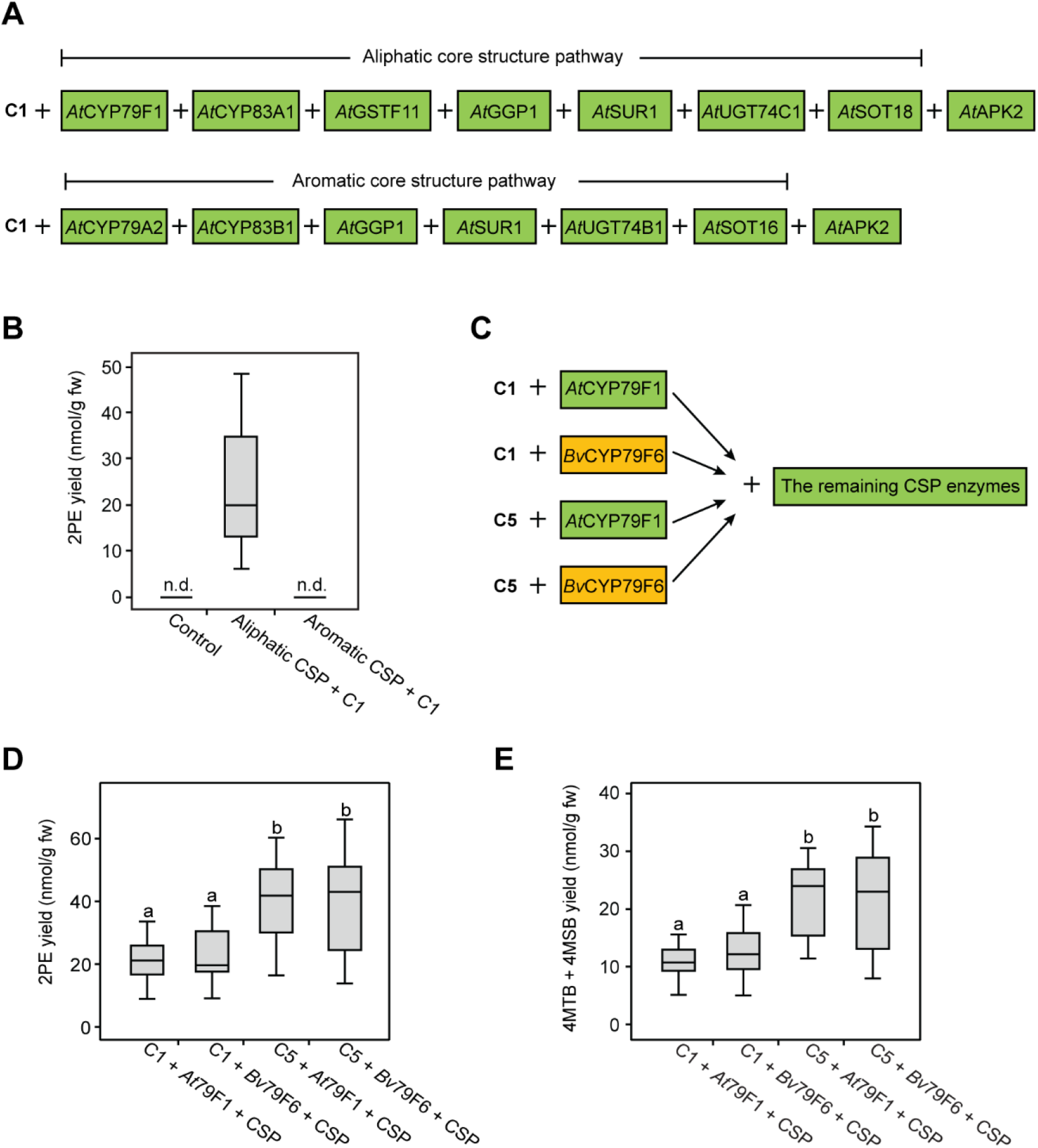
Engineering and optimization of 2PE production in *N. benthamiana* upon transient expression of different pathway combinations. **A)** Combinations of enzymes including the chain elongation combination C1, the aliphatic or aromatic core structure pathways as well as the assisting enzyme APK2 from *A. thaliana*. **B)** 2PE production by co-expression of C1 with the aromatic and aliphatic core structure pathways, respectively. “Aliphatic CSP + C1” represents the combination containing enzymes of the chain elongation combination C1, the aliphatic core structure pathway and *At*APK2. “Aromatic CSP + C1” represents the combination containing enzymes of the chain elongation combination C1, aromatic core structure pathway and *At*APK2. Control plants were infiltrated with p19 and empty plasmid. **C)** Schematic representation of the tested biosynthetic pathway combinations including chain elongation combination C1 or C5 together with the entry point either *At*CYP79F1 or *Bv*CYP79F6 (substituting the initial CYP79F shown in **A**) plus the remaining enzymes shown in **A**. **D)** 2PE production by expression of the combinations given in **C**. **E)** Production of 4MTB and 4MSB by expression of the combinations given in **C**. In **B**, data of each box represent nine biological replicates in nanomole per gram fresh weight. In **D** and **E**, data of each box represent 16 biological replicates. Data in **B**, **D** and **E** are box-and- whisker representations indicating the 10th (lower whisker), 25th (base of box), 75th (top of box) and 90th (top whisker) percentiles. The line within the box is median and outliers are not shown. Exact values are listed in **Supplementary Table S3**. n.d. represents not detected. Letters represent significant differences by pairwise comparisons based on Tukey HSD test (*p* < 0.05) and *p*-values are listed in **Supplementary Table S4**. Data labelled with different letters are significantly different and the labels containing the same letter mean no significant difference.

Towards further improving 2PE production in tobacco, we investigated whether the ortholog of *CYP79F1* (called *CYP79F6*) from *B. vulgaris* would be a better entry point for the core structure pathway. We used the reported sequence of *CYP79F6* from the *B. vulgaris* transcriptome dataset (Liu et al., 2016) that shares 83% similarity with *At*CYP79F1. No significant difference was observed in production of HPhe and DHM-derived GLSs when replacing *At*CYP79F1 with *Bv*CYP79F6 (**Fig. 3D** and **E**), showing that the two CYP79F enzymes are equally efficient in converting HPhe or DHM to the corresponding oximes in tobacco. Furthermore, co-expression of the chimeric C5 with the core structure pathway containing *Bv*CYP79F6 increased the production of HPhe and DHM-derived GLSs by around 2-fold compared to when C1 was co-expressed (**Fig. 3D** and **E**). This tendency further suggests that the chimeric C5 displays a higher flux of chain elongation machinery than the *B. vulgaris* combination C1 for GLS biosynthesis. Additionally, the accumulation of HM-derived 3MSP was maintained at low level (around 0.2 nmol/g fw) from the four pathway combinations (**Supplementary Fig. S4**), corresponding to the production level of HM by only expression of C1 and C5 (0.13 and 0.12 nmol/g fw, respectively). This result shows that the differences between the pathway combinations do not affect the accumulation level of HM-derived 3MSP.

### 3.5. The levels of precursor and chain-elongated amino acids are not reflected in the GLS profile

To investigate the effect of the different pathway combinations on the levels of precursor amino acids, we analyzed the levels of phenylalanine and methionine in each situation. Expression of only the chain elongation pathways affected neither the phenylalanine nor the methionine levels (**Fig. 4A**). The large difference between pool size of the phenylalanine (94, 97 and 80 nmol/g fw) and methionine (13, 14 and 11 nmol/g fw) in, respectively, control, C1- and C5-expressing tobacco was not reflected in the levels of HPhe and DHM as comparable levels were produced (15 and 20 nmol/g fw versus 24 and 20 nmol/g fw for C1 and C5, respectively) (**Supplementary Table S3**). Addition of the core structure pathways led to a significant reduction of phenylalanine levels, but not methionine levels compared to the control (**Fig. 4A**), suggesting that the methionine pool is more efficiently replenished than the phenylalanine pool in tobacco. The increase of 2PE production with chain elongation combination C5 compared to C1, along with the core structure pathway, did not lead to a distinguishable further lowering of phenylalanine levels, suggesting that the required higher flux is sustained by increased phenylalanine biosynthesis (**Fig. 4A**).

**Fig. 4.**
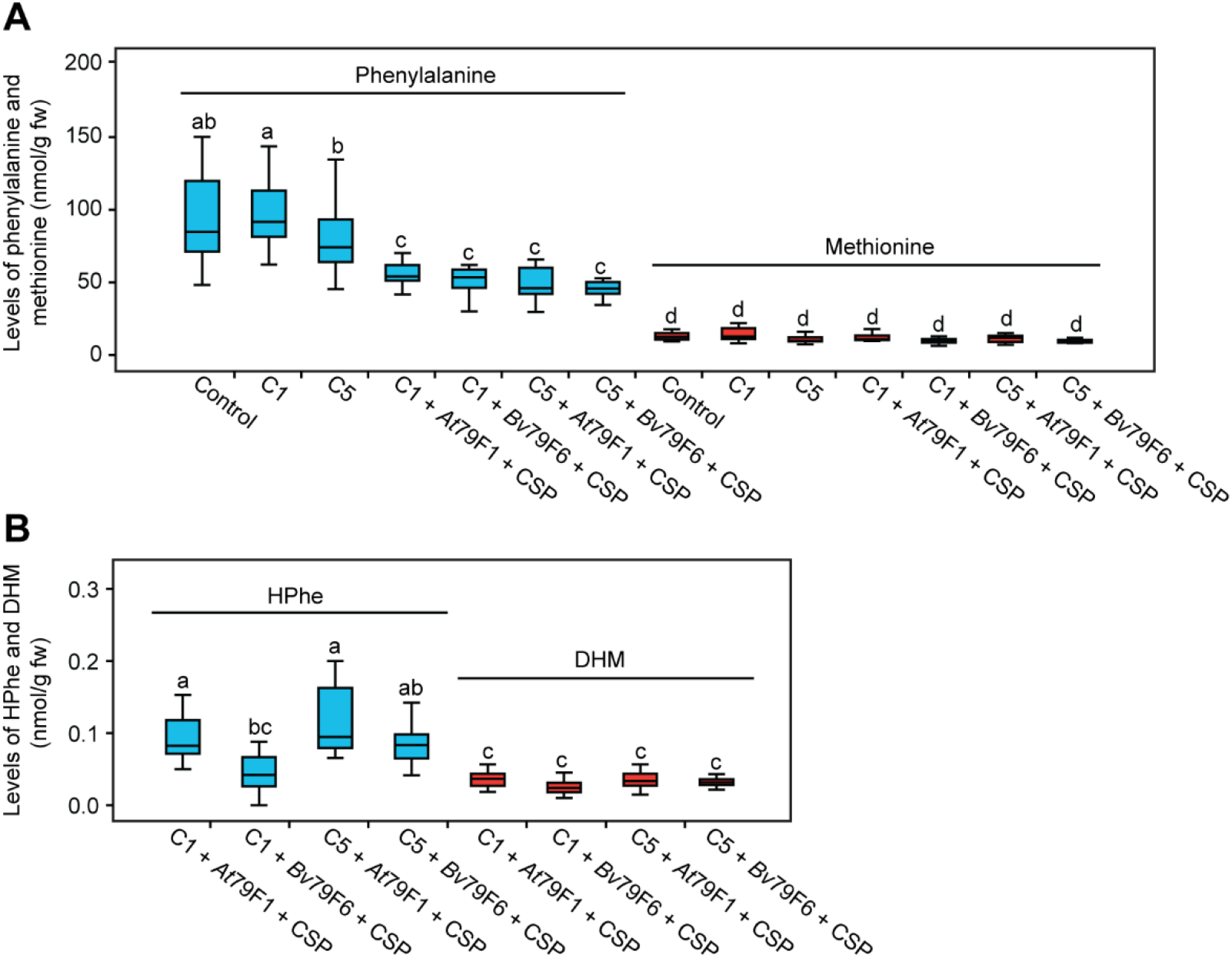
Levels of phenylalanine, methionine, HPhe and DHM in *N. benthamiana* upon transient expression of different pathway combinations. **A)** Levels of the precursors phenylalanine and methionine when expressing the different pathway combinations. The blue and red boxes indicate the levels of phenylalanine and methionine, respectively. Control plants were infiltrated with p19 and empty plasmid. Abbreviations for explanation of combinations are given in **Fig. 1A** and **Fig. 3C**. **B)** Levels of the intermediates HPhe and DHM by expression of the entire biosynthetic pathway combinations. The blue and red boxes indicate the levels of HPhe and DHM, respectively. Data of each box represent 16 biological replicates in nanomole per gram fresh weight. Data are box-and-whisker representations indicating the 10th (lower whisker), 25th (base of box), 75th (top of box) and 90th (top whisker) percentiles. The line within the box is median and outliers are not shown. Exact values are listed in **Supplementary Table S3**. Letters represent significant differences by pairwise comparisons based on Tukey HSD test (*p* < 0.05) and *p*-values are listed in **Supplementary Table S4**. Data labelled with different letters are significantly different and the labels containing the same letter mean no significant difference.

To investigate the efficiency of the coupling between chain elongation and core structure pathway, we analyzed the levels of chain-elongated amino acids as intermediates linking chain elongation and core structure pathway. The levels of HPhe and DHM were reduced to around 0.1 nmol/g fw when the core structure pathway was co-expressed (**Fig. 4B** and **Supplementary Table S3**), which shows that a coupling between the chain elongation and core structure pathway is established. Furthermore, we found decreased levels of HM, THM, HL and DHL compared to levels when the chain elongation pathways were expressed alone - especially THM reached trace level (**Supplementary Fig. S5**). However, the corresponding GLSs except HM-derived 3MSP were not detected, which suggests that the core structure pathway was not completed or that the chain-elongated amino acids are converted to other metabolites.

## 4. Discussion

In this study, we engineered 2PE production by co-expressing the proposed *B. vulgaris* enzymes in the phenylalanine chain elongation pathway and the *A. thaliana* aliphatic core structure pathway in tobacco. We show that the first two enzymes *Bv*BCAT4 and *Bv*MAM1 in the chain elongation pathway are critical for high production of HPhe, and that 2PE production can be increased 2-fold by co-expressing a chimeric combination of the chain elongation pathway (*Bv*BCAT4, *Bv*MAM1, *At*IPMI and *At*IPMDH1) compared to the chain elongation pathway from *B. vulgaris*.

The enzymes of *B. vulgaris* and *A. thaliana* are qualitatively redundant in chain elongation of phenylalanine, methionine and leucine when heterologously expressed in tobacco (**Fig. 1** and **Supplementary Fig. S2**), although the quantitative product profiles differ. In contrast, the investigated P- and G-types of *B. vulgaris* do not accumulate GLSs derived from chain-elongated aliphatic amino acids (Agerbirk et al., 2015), and wild type *A. thaliana* does not accumulate leucine-derived GLSs (Brown et al., 2003; Hanschen et al., 2018). This contrast between heterologous and native products of enzymes suggests that in the native plants, some amino acid pools are spatially separated from the site of chain elongation or core structure biosynthesis, or that additional proteins affect realized specificity *in planta*. The contrast was expectable since the chain elongation enzymes from *A. thaliana* have also previously shown activity towards leucine when expressed in *N. benthamiana* and *E. coli* (Mikkelsen et al., 2010; Mirza et al., 2016).

Surprisingly, the *B. vulgaris* enzymes produce around the same level of DHM and THM as of HPhe when expressed in tobacco, suggesting that substrate specificity of the investigated *B. vulgaris* enzymes with respect to the methionine side chain is not the restriction in native *B. vulgaris*, where methionine-derived GLSs are lacking. An obvious alternative explanation for such GLS profile *in planta* is the lack of transport of the first (α-keto acid) intermediate from cytoplasm to chloroplast, but other factors may also influence the combined product profile of chain elongation.

From an engineering point of view, it was significant that the *Bv*BCAT4 enzyme is quantitatively better than *At*BCAT4 as entry enzyme, for both the production of HPhe and DHM. Thus, the chimeric combination C4 with *Bv*BCAT4 and the remaining *A. thaliana* enzymes seems a better choice than the *A. thaliana* combination C3 for production of 4MSB. The second chain elongation enzyme, MAM1, plays a critical role in the profile of chain-elongated amino acids (**Fig. 1**). Previously, *At*MAM1 was shown to be responsible for HPhe production in *A. thaliana*, and heterologous expression of *At*MAM1 in *E. coli* enables HPhe production (Petersen et al., 2019b). Likewise, a MAM1 homolog in *Brassica napus* has been genetically linked to 2PE formation (Kittipol et al., 2019). Our data suggests that *Bv*MAM1 has substrate specificity towards α-keto acids from both phenylalanine and methionine. However, *At*MAM1 appears to prefer the α-keto acid from methionine to the α-keto acid from phenylalanine (**Fig. 1**), consistent with the fact that 2PE is a minor GLS in *A. thaliana* (Brown et al., 2003). The recent crystal structure of MAM1 enzyme from *Brassica juncea* (Kumar et al., 2019) may contribute to identify the residues responsible for the substrate specificity of *Bv*MAM1 and *At*MAM1. For instance, *At*MAM1 mutations in the residues Thr257 and Gly259 have previously been shown to increase or decrease HPhe levels in *E. coli* (Petersen et al., 2019b). Despite the broad substrate specificity of *Bv*MAM1, it is noteworthy that the inclusion of this enzyme in C5 (compared to having *At*MAM1 in C4) changed the product profile from the *A. thaliana* type to the *B. vulgaris* type. This profile shift demonstrates a reduced tendency of *Bv*MAM1 to contribute to methionine chain elongation compared to the *At*MAM1, in agreement with the native *B. vulgaris* GLS profile (**Fig. 1**).

Noticeably, in our study the CYP79 enzymes metabolizing HPhe belong to the CYP79F subfamily and not to the CYP79A subfamily, which otherwise metabolizes aromatic amino acids (Wittstock and Halkier, 2000; Koch et al., 1995). The CYP79F enzymes was previously shown to solely accept aliphatic chain-elongated derivatives of methionine and leucine as substrate (Chen et al., 2003; Mikkelsen et al., 2010). Therefore, our result indicates that the substrate specificities of CYP79F enzymes are not limited to aliphatic but also include aromatic amino acids, thereby resembling other CYP79s such as CYP79Cs and CYP79Ds (Wang et al., 2020). We anticipated that engineering with *Bv*CYP79F6 would boost 2PE production, but our study shows that *Bv*CYP79F6 and *At*CYP79F1 resulted in the same level of 2PE (**Fig. 3D**). Hence, the initial enzyme CYP79F in the core structure pathway seems not to be critical for the combined product profile of *B. vulgaris* GLS biosynthesis. Collectively, our results suggest that the 2PE biosynthesis in *B. vulgaris* is based on orthologs of the biosynthesis of methionine-derived GLS, but with specificity favoring 2PE. This finding helps to explain the enigmatic evolutionary transition from methionine to phenylalanine-derived GLSs in *B. vulgaris* (Byrne et al., 2017) and the intermediate phenotype of closely related *N. officinale* (Agerbirk et al., 2014).

Although the C1 and C5 combination of chain elongation pathways produced similar levels of HPhe when they were expressed alone (**Fig. 1B**), 2PE levels increased 2-fold when the core structure pathway was co-expressed with the chimeric chain elongation pathway C5 compared to C1 consisting of *B. vulgaris* genes only (**Fig. 3D**). Interestingly, the same situation existed for production of DHM-derived 4MTB and 4MSB (**Fig. 1C** and **Fig. 3E**). The only differences between these two biosynthetic pathways are IPMI (consisting of LSU1 and SSU3) and IPMDH1, the product of which is transaminated by BCAT4 to form the substrate of the CYP79F enzyme (**Fig. 5**). It shows that the flux of the chimeric chain elongation pathway C5 is promoted more than the all *B. vulgaris* enzymes C1 by the presence of the core structure pathway, and suggests that the increased yield of construct C5 is due to the higher *in planta* capacity of the *At*IPMI and *At*IPMDH1 enzymes than the corresponding *B. vulgaris* enzymes. Indeed, in pathway engineering, natural biodiversity is often exploited to produce and/or optimize production of valuable compounds. For example, a biosynthetic pathway for rosmarinic acid was improved by using rosmarinic acid synthases from three different plant species in *E. coli* (Bloch and Schmidt-Dannert, 2014), and a chimeric biosynthetic pathway containing enzymes from three different organisms boosted *n*-butanol production in *E. coli* (Bond-Watts et al., 2011).

**Fig. 5.**
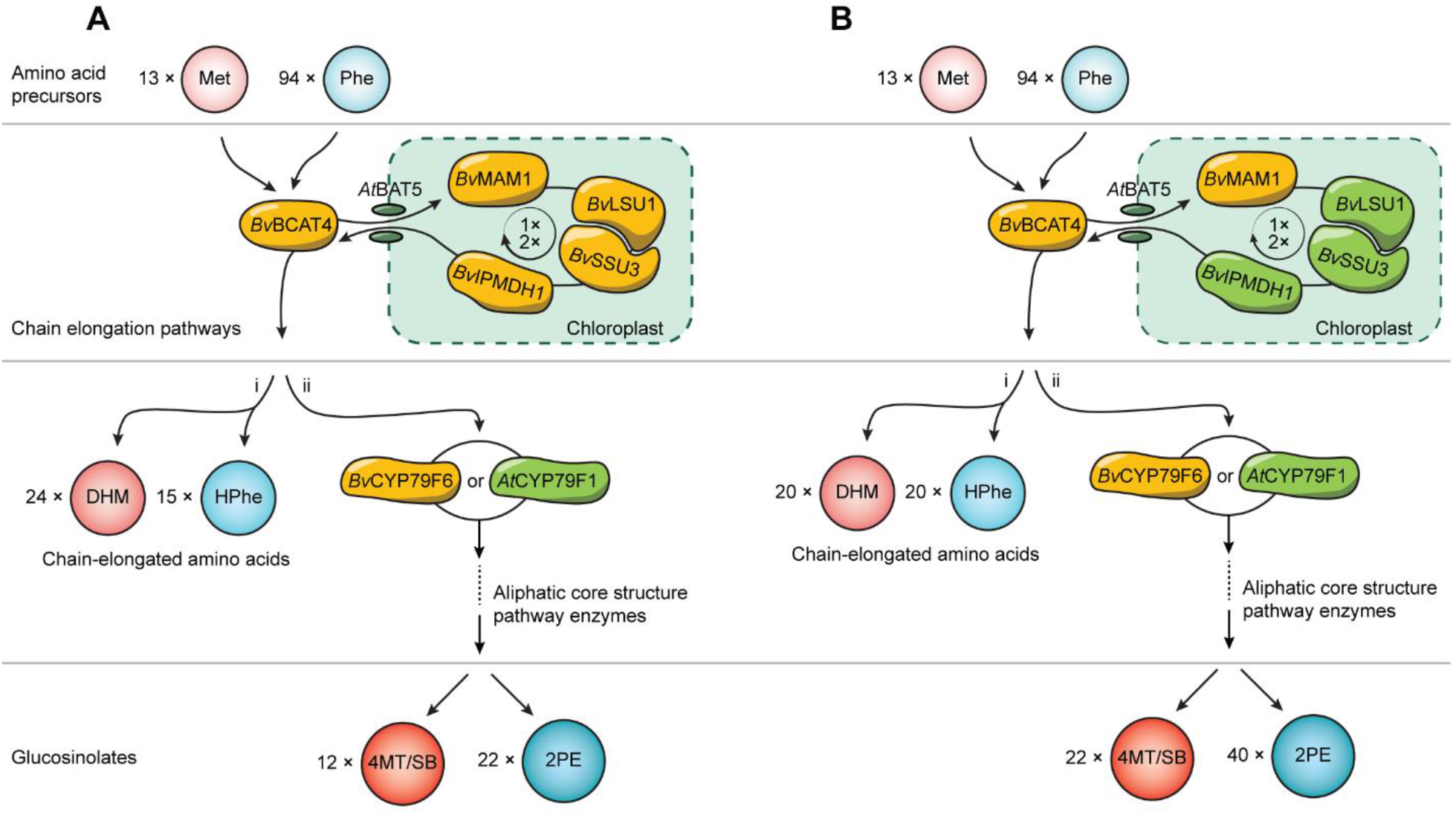
Overview of pool sizes of amino acid precursors and production levels of chain-elongated and the corresponding GLS end products for the engineered GLS biosynthetic pathways. **A)** The C1 chain elongation pathway (i) and the C1 chain elongation pathway + CYP79Fs + remaining aliphatic core structure pathway (ii). **B)** The C5 chain elongation pathway (i) and the C5 chain elongation pathway + CYP79Fs + remaining aliphatic core structure pathway (ii). LSU1 and SSU3 are the two subunits of IPMI enzyme. The light red and light blue spheres represent methionine (Met) and phenylalanine (Phe), respectively. The red and blue spheres represent dihomomethionine (DHM) and homophenylalanine (HPhe), respectively. The dark red sphere represents the sum of 4-(methylthio)butylglucosinolate (4MTB) and 4-(methylsulfinyl)butylglucosinolate (4MSB) and the dark blue sphere represents 2-phenylethylglucosinolate (2PE). The number of spheres indicates the relative molar amounts of corresponding compounds.

During engineering the 2PE production, we also detected DHM-derived 4MTB and 4MSB from the same pathway, yet with the total level as half as 2PE (**Fig. 5**). The higher yield of 2PE could be reflected by a higher substrate availability of phenylalanine compared to methionine (**Fig. 4A**). This hypothesis, however, would be in contrast to the statistically insignificant tendency for the chain elongation pathway (C1) to produce more DHM than HPhe (*p* = 0.10) despite much lower level of methionine than phenylalanine (**Supplementary Table S3**). Additionally, our result suggests that the core structure pathway containing *At*SOT18 from ecotype Col-0 efficiently catalyzes 2PE formation, which is opposite to the previous finding that *At*SOT18 exhibits no activity towards desulfo-2PE (Klein et al., 2006). An explanation for this observation could be that an endogenous tobacco SOT enzyme rather than the introduced *At*SOT18 catalyzes conversion of desulfo-2PE to 2PE.

HPhe is an intermediate in 2PE biosynthesis, but also a versatile pharmaceutical precursor for medicines (Brown and Vaughan, 1998; Kuhn et al., 2007). Thus, researchers have attempted production of HPhe in a sustainable and efficient way. For example, HPhe was synthesized using the substrates phenylalanine and α-keto-4-phenylbutyric acid by an aromatic aminotransferase in yeast (Hwang et al., 2009). Three putative genes were identified from the cyanobacterium *Nostoc punctiforme* and co-expressed in *E. coli* to achieve HPhe production (Koketsu et al., 2013). Increased production of HPhe was observed by expressing an engineered NADPH-dependent glutamate dehydrogenase in *E. coli* (Li and Liao, 2014). In our study, the engineering of the chimeric chain elongation pathway C5 provides a new approach to produce HPhe for medicinal application.

In conclusion, we have engineered the production of HPhe and 2PE in *N. benthamiana*. This achievement has great potential for stably integrating the biosynthesis of 2PE to plants and microorganisms for the benefit of human health, defensive properties of crops, and tailoring of beneficial microbes.

## Abbreviations

GLS: glucosinolate;
2PE: 2-phenylethylglucosinolate;
HPhe: homophenylalanine;
HM: homomethionine;
DHM: dihomomethionine;
THM: trihomomethionine;
HL: homoleucine;
DHL: dihomoleucine;
PEITC: 2-phenylethyl isothiocyanate;
3MTP: 3-(methylthio)propylglucosinolate;
3MSP: 3-(methylsulfinyl)propylglucosinolate;
4MTB: 4-(methylthio)butylglucosinolate;
4MSB: 4-(methylsulfinyl)butylglucosinolate;
BGLS: benzylglucosinolate

## Author contributions

CW, CC and BAH designed the experiments. CW performed all the experiments and CC performed LC-MS method development and analysis. CW, CC, NA and BAH interpreted the data and wrote the manuscript based on a draft by CW.

## Acknowledgements

This work was supported by the Danish National Research Foundation [grant DNRF99] and the Novo Nordisk Foundation [NNF17OC0027710]. Mads Møller Dissing is thanked for helping to clone the constructs for expressing *Bv*BCAT4 and *Bv*MAM1 and Professor emerita M. Soledade C. Pedras is thanked for advice on organic nomenclature.

