## Supplementary Fig. S1-S5 for "Engineering and optimization of the 2-phenylethylglucosinolate production in *Nicotiana benthamiana* by combining biosynthetic genes from *Barbarea vulgaris* and *Arabidopsis thaliana*"

**Contents:**

**Supplementary Fig. S1.** The proposed biosynthetic pathway of 2PE.

**Supplementary Fig. S2.** Production of HM, THM, HL and DHL in *N. benthamiana* upon transient expression of the different chain elongation combinations.

**Supplementary Fig. S3.** Production of 3MSP, 4MTB, 4MSB and BGLS upon co-expression of the chain elongation combination C1 together with respectively aliphatic or aromatic core structure pathway in *N. benthamiana*.

**Supplementary Fig. S4.** Production of 3MSP in *N. benthamiana* upon transient expression of the different pathway combinations.

**Supplementary Fig. S5.** The levels of HM, THM, HL and DHL in *N. benthamiana* upon transient expression of the different pathway combinations.

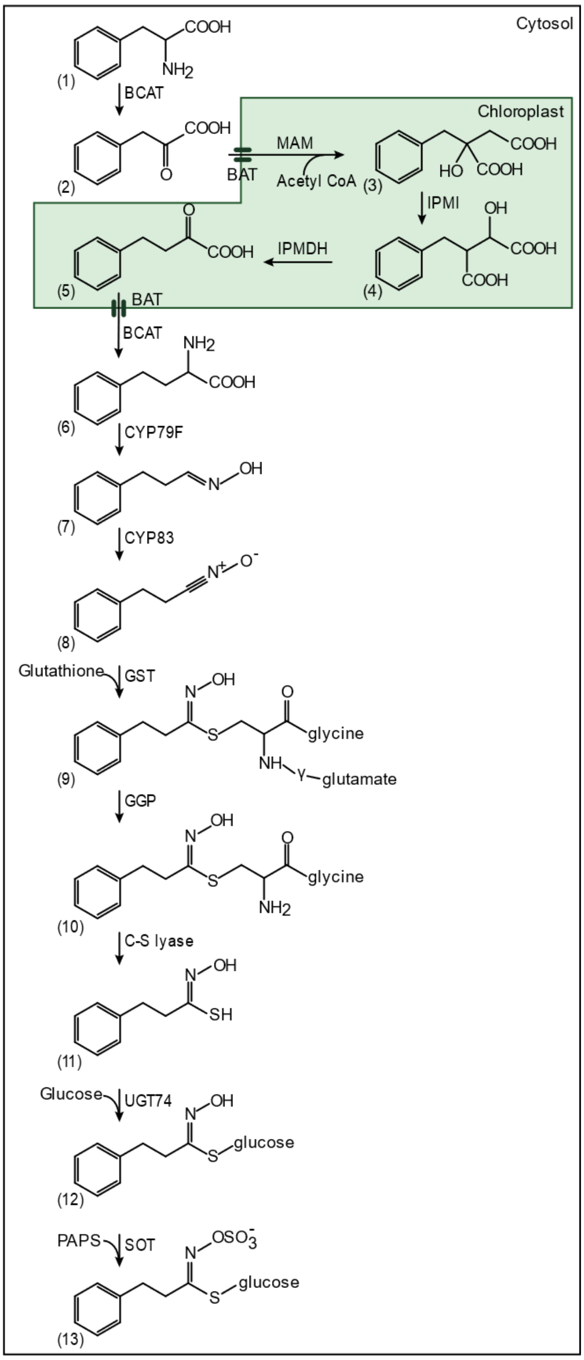


**Supplementary Fig. S1. The proposed biosynthetic pathway of 2PE.** The green box represents the chloroplast, in which three of the chain elongation pathway reactions occur. The remaining part represents the cytosol, with the first reaction of chain elongation as well as the core structure pathway taking place after chain elongation. The enzymes BCAT, BAT, MAM, IPMI, IPMDH, CYP79F, CYP83, GST, GGP, UGT74 and SOT are, respectively, branched-chain amino acid aminotransferase, bile acid transporter, methylthioalkylmalate synthase, isopropylmalate isomerase, isopropylmalate dehydrogenase, cytochrome P450 of the CYP79F subfamily, cytochrome P450 of the CYP83 family, glutathione-*S*-transferase, γ-glutamate peptidase, UDP-glucosyltransferase of the 74 family and sulfotransferase. The compounds in the pathway are phenylalanine (1), 3-phenyl-2-oxopropanoic acid (2), 2-benzyl-2-hydroxysuccinic acid (3), 2-benzyl-1-hydroxysuccinic acid (4), 4-phenyl-2-oxobutanoic acid (5), homophenylalanine (6), (*E*)-3-phenylpropanal oxime (7), 3-phenylpropanenitrile *N*-oxide (8), *S*-[(*Z*)-3-phenylpropylhydroximoyl]-L-glutathione (9), *S*-[(*Z*)-3-phenylpropylhydroximoyl]-L-Cys-L-Gly (10), (*Z*)-3-phenylpropanthiohydroximic acid (11), desulfo-2-phenylethylglucosinolate (12), 2-phenylethylglucosinolate (13).

**Supplementary Fig. S2. Production of HM, THM, HL and DHL in *N. benthamiana* upon transient expression of the different chain elongation combinations. A)** HM yields. **B)** THM yields. **C)** HL yields. **D)** DHL yields. Data of each box represent 16 biological replicates in nanomole per gram fresh weight. Data in **A**-**D** are box-and-whisker representations indicating the 10th (lower whisker), 25th (base of box), 75th (top of box) and 90th (top whisker) percentiles. The line within the box is median and outliers are not shown. Exact values are listed in **Supplementary Table S3**. n.d. represents not detected. Letters represent significant differences by pairwise comparisons based on Tukey HSD test ($p < 0.05$) and $p$-values are listed in **Supplementary Table S4**. Data labelled with different letters are significantly different and the labels containing the same letter mean no significant difference.

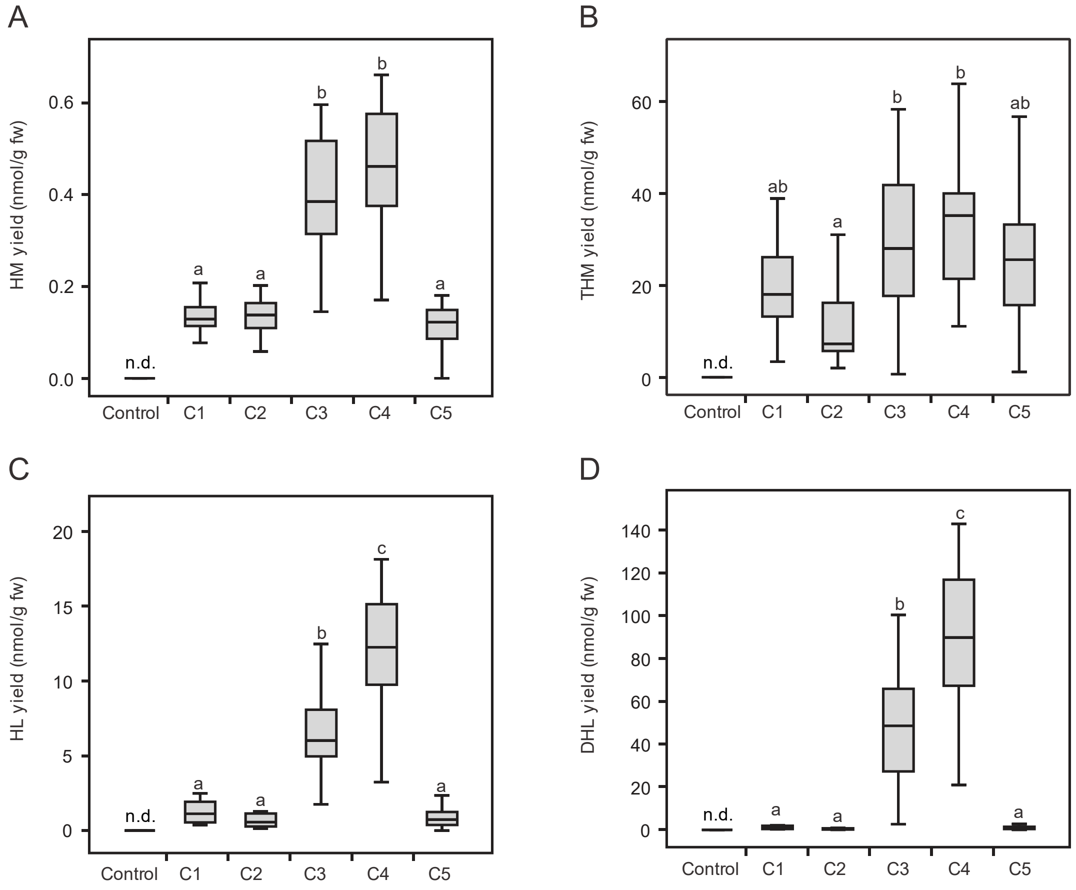

**Supplementary Fig. S3. Production of 3MSP, 4MTB, 4MSB and BGLS upon co-expression of the chain elongation combination C1 together with respectively aliphatic or aromatic core structure pathway in *N. benthamiana*. A)** GLSs accumulated with co-expression of the chain elongation combination C1 and the aliphatic core structure pathway. **B)** GLS accumulated with co-expression of the chain elongation combination C1 and the aromatic core structure pathway. Data of each box represent nine biological replicates in nanomole per gram fresh weight. The box-and-whisker representations indicate the 10th (lower whisker), 25th (base of box), 75th (top of box) and 90th (top whisker) percentiles. The line within the box is median and outliers are not shown. Exact values are listed in **Supplementary Table S3**. n.d. represents not detected.

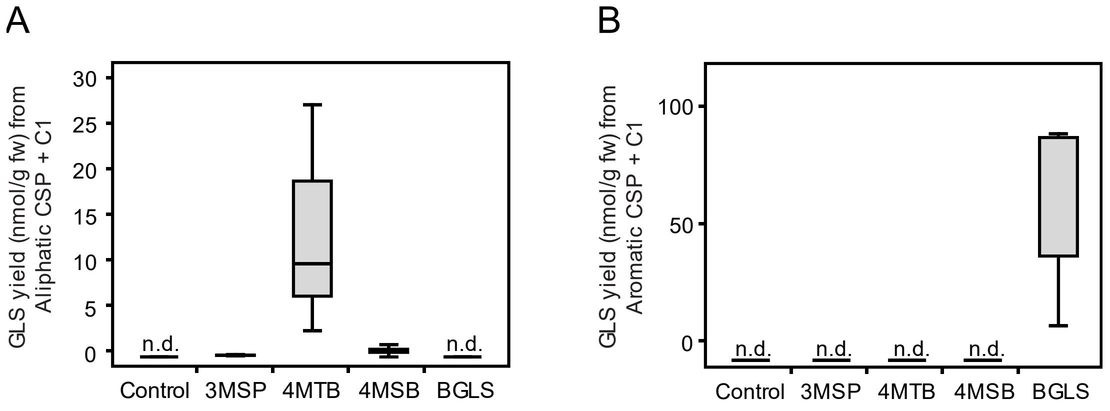

**Supplementary Fig. S4. Production of 3MSP in *N. benthamiana* upon transient expression of the different pathway combinations.** Abbreviations for explanation of combinations are given in **Fig. 3C**. Data of each box represent 16 biological replicates in nanomole per gram fresh weight. The box-and-whisker representations indicate the 10th (lower whisker), 25th (base of box), 75th (top of box) and 90th (top whisker) percentiles. The line within the box is median and outliers are not shown. Exact values are listed in **Supplementary Table S3**. n.d. represents not detected. Letters represent significant differences by pairwise comparisons based on Tukey HSD test ($p < 0.05$) and $p$-values are listed in **Supplementary Table S4**. Data labelled with different letters are significantly different and the labels containing the same letter mean no significant difference.

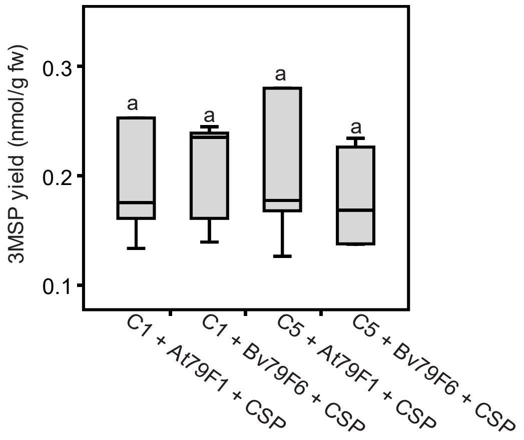

**Supplementary Fig. S5. The levels of HM, THM, HL and DHL in *N. benthamiana* upon transient expression of the different pathway combinations. A)** HM level. **B)** THM level. **C)** HL level. **D)** DHL level. Abbreviations for explanation of combinations are given in **Fig. 1A** and **Fig. 3C**. Data of each box represent 16 biological replicates in nanomole per gram fresh weight. Data in **A**-**D** are box-and-whisker representations indicating the 10th (lower whisker), 25th (base of box), 75th (top of box) and 90th (top whisker) percentiles. The line within the box is median and outliers are not shown. Exact values are listed in **Supplementary Table S3**. n.d. represents not detected. Letters represent significant differences by pairwise comparisons based on Tukey HSD test ($p < 0.05$) and $p$-values are listed in **Supplementary Table S4**. Data labelled with different letters are significantly different and the labels containing the same letter mean no significant difference.

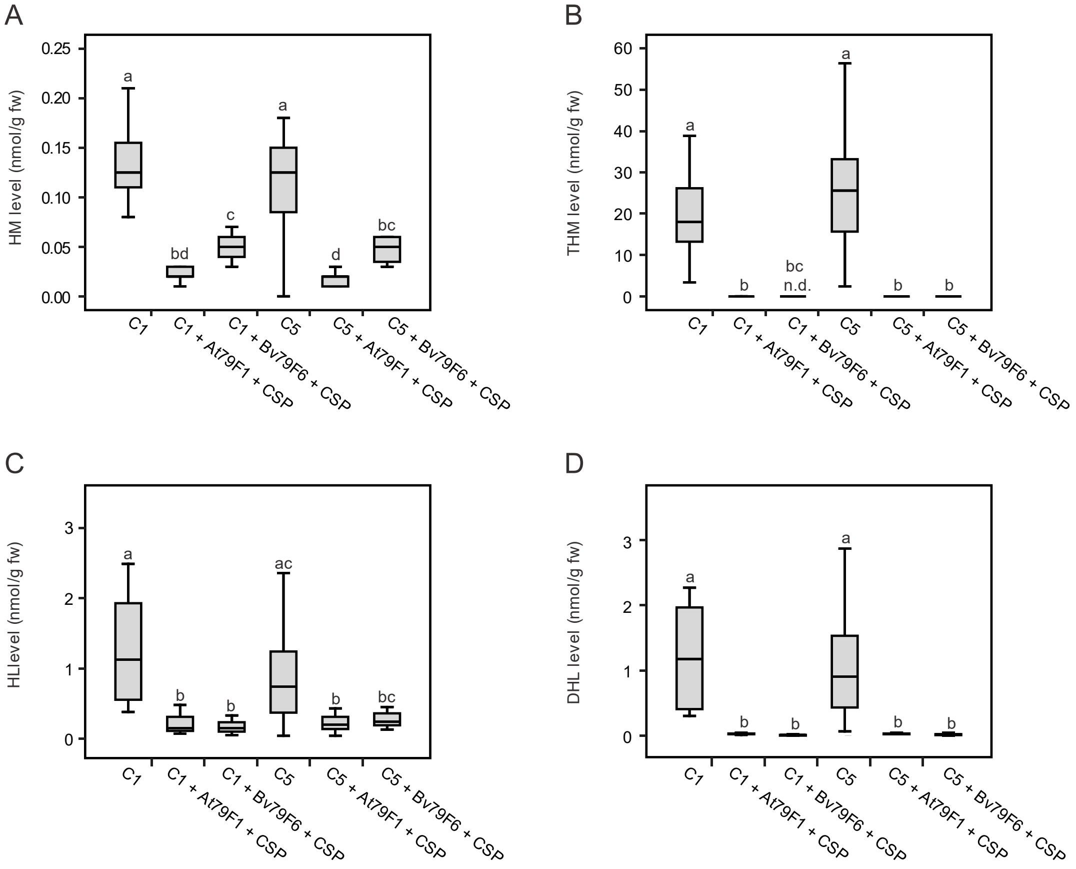
